# Effects of genomic and functional diversity on stand-level productivity and performance of non-native Arabidopsis

**DOI:** 10.1101/2020.04.20.049593

**Authors:** Kathryn G. Turner, Claire M. Lorts, Asnake T. Haile, Jesse R. Lasky

## Abstract

Biodiversity can affect the properties of groups of organisms, such as ecosystem function and the persistence of colonizing populations. Genomic data offer a newly available window to diversity, complementary to other measures like taxonomic or phenotypic diversity. We tested whether native genetic diversity in field experimental stands of *Arabidopsis thaliana* affected their aboveground biomass and fecundity in their colonized range. We constructed some stands of genotypes that we *a priori* predicted would differ in performance or show overyielding. We found no relationship between genetic diversity and stand total biomass. However, increasing stand genetic diversity increased fecundity in high resource conditions. Polyculture (multiple genotype) stands consistently yielded less biomass than expected based on the yields of component genotypes in monoculture. This under-yielding was strongest in stands with late-flowering and high biomass genotypes, potentially due to interference competition by these genotypes. Using a new implementation of association mapping, we identified genetic loci whose diversity was associated with stand-level yield, revealing a major flowering time locus associated with under-yielding of polycultures. Our field experiment supports community ecology studies that find a range of diversity-function relationships. Nevertheless, our results suggest diversity in colonizing propagule pools can enhance population fitness. Furthermore, interference competition among genotypes differing in flowering time might limit the advantages of polyculture.

## INTRODUCTION

The characteristics of groups of organisms can affect their persistence and collective functions. The emergent properties of groups of organisms are major topics across biology [1–3]. The mechanisms underlying the benefits of diversity for group-level performance can be divided into at least two categories [4]. First, greater diversity can reduce the negative effects associated with high density populations or communities. This can occur via diversity in resource use that reduces competition, diversity in susceptibility to enemies that reduces “apparent” competition, or facilitation among diverse individuals. Second, diversity can increase group-level performance via a sampling effect: greater diversity increases the chance that high-performing individuals will be present. Diversity can also reduce group-level performance. This can occur when more diverse groups tend to have lower performing individuals (e.g. when diversity is driven by immigration of maladapted individuals) [5,6] or when diversity results in interference competition among different types [7]. While diversity effects on ecosystem functions (EF) can act at both population and community levels, they have been mostly studied in communities [8,9]. However, *intraspecific* diversity can have major impacts on EF, potentially surpassing interspecific effects when a few species dominate biomass [10].

Community ecologists have often hypothesized that diversity increases ecosystem functions [11,12] while invasion biologists have hypothesized that diversity increases the growth of colonizing populations [13–15]. These fields have been largely independent and disjunct, but they share much in common (see also related topics in kin and group selection [16]). Both address hypotheses based on the same underlying biological processes: 1) sampling effects, whereby greater diversity increases the chance a group will contain high functioning/performing individuals, 2) insurance effects (a type of sampling effect), whereby greater diversity increases the chance that a group will contain an individual that is high functioning under a new environment, or 3) complementarity effects, whereby diversity relieves negative density effects. Invasion biology has long investigated how the number of individuals in a founding group affects colonization success [17,18]. Evidence suggests that increased genetic diversity, rather than increased numbers of individuals *per se*, causes improved establishment [13], growth rate [13–15], biomass production [2,19,20], and persistence of colonizing populations, and decreases vulnerability to environmental change [21]. The effects on group-level performance and EF are likely to be conditional on the environment. For example, facilitation may dominate in harsh environments, strengthening positive diversity effects [22,23], or novel environments might be more limited by diversity due to a rarity of high-functioning individuals. Empirical studies of genetic diversity in invasions have found it favorable under some conditions but not others [2,24] and theoretical predictions also conflict [20,25].

Diversity effects on the performance of a group of organisms arise fundamentally due to the effects of phenotypic diversity [26]. However, the phenotypes that affect group-level performance are very often unknown. Instead, genetic, taxonomic, or phylogenetic diversity can be useful proxies. Past studies of intraspecific diversity and EF have used either genotypes of unknown relatedness [19,20,27], or have used a small number of genetic markers having unclear connections to functional variation [2,28,29]. By contrast, large numbers of markers across the genome allow precise estimation of genetic similarity between individuals, and it is now feasible to measure whole genome diversity in many non-model systems. Furthermore, genetic diversity effects may be most strongly associated with the specific loci that underly complex phenotypes associated with resource economics. Genetic variation in complex traits corresponding to life history and resource use likely involves changes in many loci [30,31], and simple genome-wide additive models can often predict variation in such traits [32–34]. Additionally, genetic mapping may be used to identify specific genes and mutations controlling phenotypes whose diversity affects EF [23,35].

Here we investigate how intraspecific diversity influences productivity and fecundity for the model plant *Arabidopsis thaliana* across environments in a field experiment outside its native range. Arabidopsis is useful for our questions: it is a model for ecological genomics [36], diverse germplasm with resquenced genomes is available [37], and it exhibits variation in resource strategies and life history [38,39] suggesting that diversity may impact EF. Our study builds upon previous work in Arabidopsis [19,23,35] by using ecologically realistic and relevant field experimental conditions, including a low resource soil treatment, and by studying stands with up to 20 genotypes. Because Arabidopsis is largely selfing and we studied a single generation, we set aside the effects of diversity on population performance due to sexual reproduction, such as heterosis.

Experimental studies of biodiversity effects typically use randomly assembled groups of genotypes or species, despite existing prior knowledge of potential drivers of compositional and diversity effects on EF. Here, in addition to randomly assembled groups representing a range of genotypic diversity, we use knowledge of our system to develop *a priori* selected groups of genotypes that we predict will exhibit particularly strong differences in EF, groups benefitting from potential niche complementarity, and groups of genotypes adapted to the experimental environment.

## METHODS

### Plant material

We studied 60 diverse Arabidopsis genotypes from across the native Eurasian range of Arabidopsis, mainly from the Mediterranean, central Asia, the UK and central Europe (Figure S1). All genotypes are representatives of natural inbred lines that had their genomes resequenced [37]. We selected these genotypes based on diversity and seed availability from a common seed production environment used in previous experiments (Methods S1).

### Field experiment

To assess the impact of different levels of genetic diversity in experimentally assembled non-native populations of Arabidopsis, we measured performance of Arabidopsis stands in a field common garden at a site (40.8106° N, 77.8472° W) in the non-native range near other established non-native populations (within ∼10 km) but with none at this site (within ∼1km). Thus our experimental stands were likely not influenced by density-associated processes driven by wild individuals nearby. To assess the environmental context dependency of diversity effects, we replicated all stand genotype compositions in two treatments (described below). Each of the 60 genotypes were grown in monoculture in three replicate stands for each of the two treatments, for a total of 360 monoculture stands (Figure S2).

We began by constructing polycultures using *a priori* hypotheses for factors likely to affect productivity. We used five different criteria to generate these ‘prediction stands’: (1) genomic diversity among accessions, (2) climate similarity of collection location to our field site, combined with genomic diversity, (3) early flowering genotypes, (4) late flowering genotypes, (5) climate similarity of collection location to our study site, combined with flowering time diversity (see details in Methods S1). Two stand compositions were chosen for each of the five prediction scenarios at each diversity level (2, 10, 20), for a total of ten prediction stand compositions for each diversity level. Each of the prediction stands were planted into both of the two treatments, for a total of 60 prediction stands.

After genotypes were assigned to prediction stands, we also generated random polycultures. Each of these polycultures were planted into both treatments with 10 replicates of diversity level each, for a total of 60 randomly assigned polyculture stands (Methods S1).

Seeds were initially sown into 20 positions within manure pots, with each position thinned to a single plant, targeting a 20-plant stand (regardless of diversity level). High resource pots were filled with potting mix and low resource pots with a high proportion of sand. Stands were germinated in growth chambers and transplanted into the field site on May 24, 2018 (Methods S1).

We measured specific leaf area (SLA) on a representative, healthy leaf from a single plant in each monoculture stand, and flowering date on all stands. We harvested each individual plant once it reached maturity. We recorded the total silique (fruit) number and the length of one representative middle silique. All remaining plants were harvested on July 18-20, as summer heat stress set in and reproduction largely ceased. We measured aboveground dry biomass on all plants. Canopy area was similarly measured for each stand just prior to the first harvested plant using Easy Leaf Area and a red 4 cm^2^ square reference [41].

### Measures of stand composition and diversity

To estimate genomic diversity of stands, we used published SNPs from resequencing data [37]. We calculated the genetic distance matrix using the R package SNPRelate [42] and then calculated the mean genetic distance among plants sown in a stand (including zero distance for individuals of the same genotype). We also estimated phenotypic variation, based on our new SLA data and published flowering time data [37]. We took breeding values of log(SLA) measured on plants in our monocultures and calculated mean for individual plants in all stands. Flowering time composition was estimated using individuals’ breeding value for the average flowering time across 10°C and 16°C in published growth chamber experiments [37].

### Statistical analyses

To gain general insight into variation in phenotypes and performance, we estimated trait plastic responses to treatment, broad-sense heritability, and genomic signature of performance in monocultures. To test for treatment effects on trait plasticity in monocultures, we implemented linear mixed models using the R package ‘VCA’ [43] and included genotype and block as random effects and soil treatments and plot x-y coordinate as fixed effects. Plastic trait responses were characterized with this soil fixed effect. We estimated broad sense heritability of stand-level traits and performance for monocultures using linear models in R where genotypes were fixed effects, taking R^2^ of these as broad sense heritability. We tested for a genomic signature in fecundity and aboveground biomass using genomic prediction models, where traits are a function of genotype-specific random effects that are correlated according to genome-wide similarity, using the R package ‘rrBLUP’ [43]. We used ten-fold cross validation to estimate the ability of genome-wide similarity to predict performance.

To address our primary research questions, we tested whether multiple measures of diversity affected stand level performance (fecundity, aboveground biomass, and leaf canopy area). In our analyses, we controlled for variation among stands in the number of plants surviving transplant or germinating late, with a minimum of 11 plants required for inclusion in analysis. We used two different counts of plants surviving transplant, on May 29 and June 12, and took the maximum, but we found results were similar when using the count on either date.

We used linear regression models to test diversity effects on stand performance while accounting for additional sources of error. The covariates were each tested in models including an interaction with the resource-limitation treatment

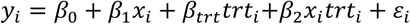

where *y*_*i*_ is the performance measure (biomass or fecundity) of stand *i* and *trt*_*i*_ is a dummy variable for resource treatment, and *x*_*i*_ is a covariate of stand *i’*s composition or diversity. Covariates *x* tested in separate models included diversity treatment level (1, 2, 10, or 20 genotypes), genomic similarity (mean genetic distance among plants in a stand), stand-level average of mean flowering time for each genotype, a measure of the stand-mean climate distance from the field site, or stand-mean SLA breeding value. We similarly tested for stand diversity and composition effects on fecundity [mean estimated total silique (fruit) length per fruiting plant in stand].

We calculated the difference between the observed polyculture biomass yields and that expected based on yields of monocultures and their proportional representation in the polyculture. This difference is a commonly studied metric in community diversity-ecosystem function studies [4], and we refer to this quantity as the deviation from monoculture expectation (DME, referred to as the “net diversity effect” by [4]). We tested how the measures of diversity and composition described above were related to DME, using linear models with a parallel structure to those used for aboveground biomass and fecundity (described above).

To determine whether our prediction stands were successful at explaining variation in fecundity and biomass production, we compared stand types using ANOVA. Different prediction criteria were coded as a factor level as were randomly assembled stands. Two-way ANOVA showed no significant treatment-stand type interaction, thus we focused on stand type effects in a two-way ANOVA without interactions. We compared each pair of stand types with post-hoc Tukey’s tests with FDR control. Because only two stands predicted to be later flowering produced fruit, we removed this plot type from our analysis of fecundity in prediction plots.

To map loci across the genome where allele frequency and variance were related to stand biomass and DME, we used an approach akin to genome wide association mapping. We tested how both stand-level per plant biomass and DME (as response variables) were associated with stand covariates of allele frequency and variance (the latter a measure of diversity for a biallelic locus), for each SNP. For a biallelic SNP, variance = *pq* where *p* and *q* are the respective frequencies of the two alleles. We used SNPs from published whole genome resequencing [37]. We excluded SNPs segregating at <0.05 minor allele frequency in our panel of 60 genotypes and across stands in the same analysis, leading to a total of ∼1.9M SNPs. For each combination of resource treatment and performance measure we tested for association with SNP allele frequency or variance. Because background variation across the genome can contribute to phenotypic variation, we calculated genome-wide similarity among stands in either allele frequency or variance at all SNPs [44]. We used a linear mixed model that included a random effect that was correlated among stands according to the genome-wide similarity matrix, using the EMMA software [44]. We also focused these association scans on SNPs within 3kb of 26 *a priori* candidate genes for flowering time [45], based on the hypothesis that genes controlling this trait might influence stand biomass and DME.

## RESULTS

### Plastic changes in traits and performance, heritability in monoculture

Comparing phenotypes for monoculture stands, SLA did not change significantly across soil treatments, nor did flowering dates (linear mixed-effects models with genotype random effects, Table S1). However, both average plant fecundity and biomass per plant in monocultures were significantly greater in the high resource treatment (Table S1, Figure 1). Date of flowering was highly heritable (∼0.8 broad-sense, depending on metric and treatment), while log(SLA), fecundity, and biomass had moderate heritability (∼0.4-0.5 depending on trait and treatment, Table S2).

**Figure 1.**
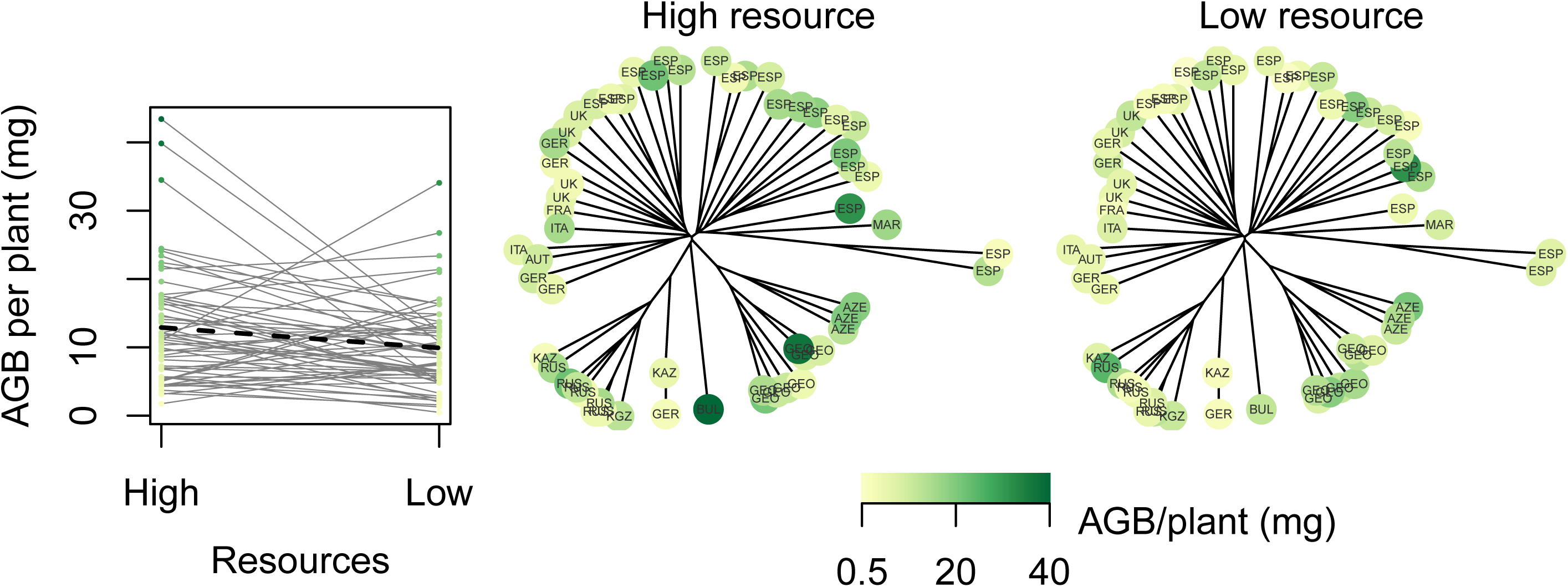
Reaction norms for aboveground biomass (AGB, left) and neighbor-joining trees, where higher yielding genotypes are colored in green. ISO country codes indicate genotype origins. Two genotypes with poor germination are not shown.

### Genomic signature of performance variation

There was a strong effect of genome-wide similarity among genotypes on biomass production in monoculture in the low resource treatment. Genomic predictions (out-of-sample) were correlated with actual breeding values for per plant biomass in monoculture in low resource (r = 0.36) but not high resource (r = −0.32) treatments. For example, in the low resource treatment a group of closely related ecotypes from the Caucasus region produced among the most biomass, while related genotypes from central Europe or Spain had low productivity. We found a similar pattern for per plant fecundity in monocultures (genomic predictions: high resource r = −0.01, low resource r = 0.32).

### How did composition and diversity of stands impact stand fecundity and biomass production?

Among all stands (including monocultures and polycultures), the low resource soil treatment significantly reduced stand fecundity, canopy coverage, and biomass production, as tested in separate linear models of each response variable (Figure 2, Tables S3-5). We found that a greater number of genotypes in a stand significantly increased the per plant fecundity in the high resource treatment (Table S3) but not the low resource treatment, while other measures of composition and diversity were unrelated with fecundity. We found that higher average breeding values for flowering time were associated with significantly greater stand biomass in both treatments, while lower breeding values for SLA was associated with greater stand biomass only in the high resource treatment (Table S4). None of the several diversity measures (i.e. the other covariates) we tested were significantly associated with biomass production.

**Figure 2.**
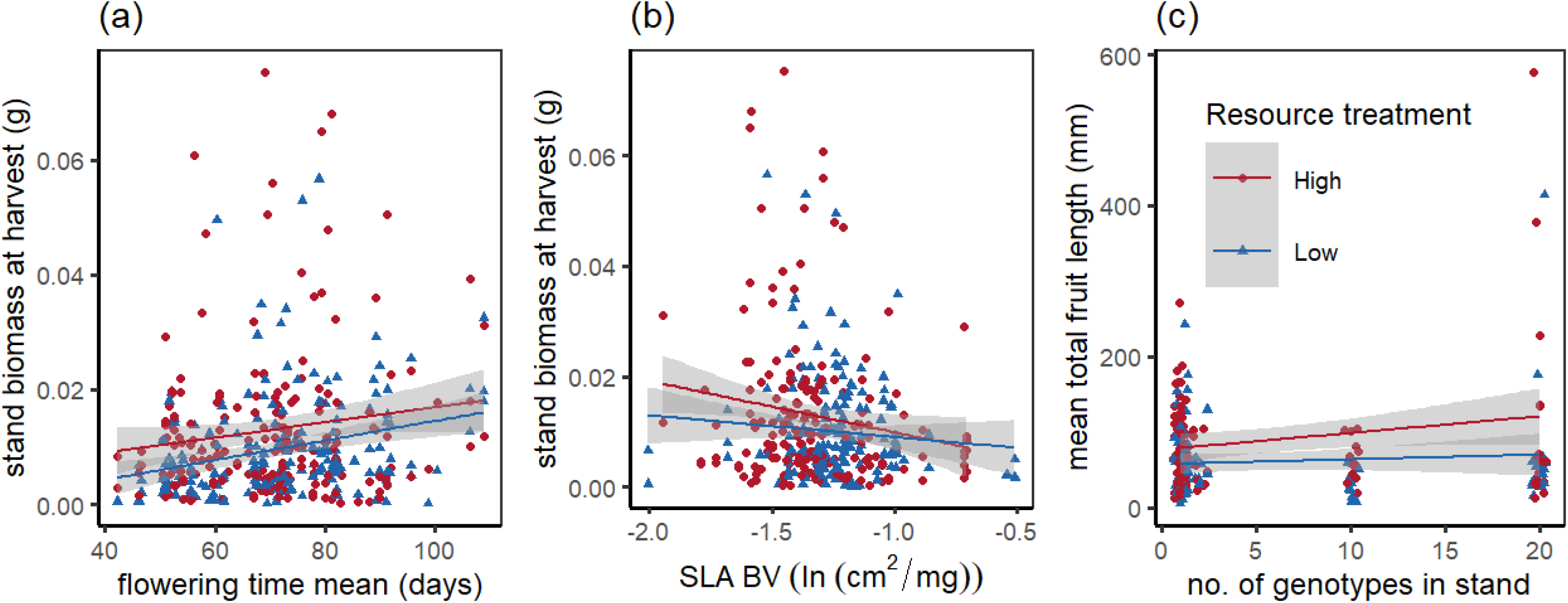
Stand-level biomass and fecundity versus composition and diversity metrics. Each point represents a stand; red circles indicate stands in the high resource treatment, blue triangles indicate the low resource treatment. Resource treatment had a significant effect on both biomass and fecundity (Tables S3-4). Shaded areas represent standard error. (a) Later stand mean breeding values for flowering time from published experiments [37] significantly increased stand biomass at harvest. (b) log(SLA) breeding value had a significant negative effect on stand biomass at harvest in the high resource treatment. (c) Experimental stand diversity level (1, 2, 10, 20 genotypes) significantly increased estimated fecundity (total fruit length) per flowering plant in the high resource treatment.

We also tested if “prediction stands” had distinct stand-level fecundity or biomass from randomly assembled stands (Figure S5). We did not find a significant effect of stand type in a two-way ANOVA for either plot level fecundity or biomass.

### How did polyculture productivity deviate from monoculture expectation?

We found that in both treatments, the yield of polyculture stands was not closely correlated with the expectation based on the yield of their component genotypes in monoculture (high resource, r = 0.051, p = 0.7138; low resource, r = 0.007, p = 0.9585). Specifically, polycultures tended to yield less than expected based on the yield of their component genotypes in monoculture. The deviation from monoculture expectation (DME) was significantly less than zero in the high resource stands (median = −61.01 mg, wilcoxon test p = 0.0025) and in the low resource stands (median = −57.13 mg, wilcoxon test p = 0.0158). These results were unchanged (both p < 0.02) when we generated randomly assembled polycultures to match the number of plants surviving transplant using genotype-specific transplant survival in monoculture.

To evaluate potential causes of negative DME, we tested correlations between DME and explanatory factors. One potential cause of negative DME was competitive inhibition of many smaller plants by late flowering plants whose rosettes grew to overtop the earlier flowering smaller plants. We found that in the high resource treatment, there was more negative DME in stands with greater average breeding values for flowering time (mean flowering time; linear model, flowering time in high resource slope = −0.0064 g/day, p = 0.0167, in low resource slope = −0.0012 g/day, p = 0.6532) or stands having plants with greater flowering time (90th percentile of flowering time; high resource slope = −0.0051 g/day, p = 0.0170, in low resource slope = −0.0006 g/day, p = 0.7883, Table S6). We also found (perhaps unsurprisingly) that biomass in monoculture of component genotypes was negatively associated with DME in polyculture: stands with greater median or 90th percentile of genotypes’ biomass in monoculture showed significantly bigger reductions in polyculture yield in both high and low resource treatments (Table S6). We tested whether DME might be due to genomic similarity, number of genotypes, or climate of origin of genotypes. We found no strong effects on DME of mean kinship, total tree diversity, number of genotypes, mean SLA, or mean distance to climate of origin along the first five PCs (Table S6).

We also tested if “prediction stands” had distinct deviations from monoculture expectation (DME). We found that there were significant differences in DME among prediction stand criteria (two-way ANOVA with treatment and stand prediction criteria, including randomly assembled stands: treatment p = 0.8675, prediction p = 0.0433, only including prediction stands: treatment p = 0.8258, prediction p = 0.0328). Consistent with the regression of DME on flowering time on all plots, the differences among our prediction plots were largely driven by differences between the predicted early (positive diversity effect) and late flowering (negative diversity effects) stands (Tukey HSD comparison of means, including randomly assembled stands, only significant difference after multiple comparisons correction is: early flowering-late flowering predicted stands DME = 0.27 g, FDR adjusted p = 0.0292).

### What genetic loci contribute to diversity effects?

We tested whether per plant biomass or DME in each treatment were associated with allele frequency or diversity at particular SNPs across the genome. We found that overall, there were few loci with dramatically stronger (outlier) associations with biomass and DME, but rather patterns suggest that the genetic basis of variation in biomass and DME were polygenic.

Nevertheless, we found that, in concordance with the association of late-flowering genotypes with under-yielding polycultures, the top SNP (chr. 2, 9584630 bp, TAIR10, genome-wide significant at FDR = 0.1, nominal p = 8.1 ×10^−7^) associated with DME in the high resource treatment (both allele frequency and variance association scans) was closest to SHORT VEGETATIVE PHASE (SVP, also known as AGL-22, AT2G22540). SVP is a MADS-box transcription factor, one of the top loci associated with natural variation in Arabidopsis flowering time in [37] (Figure 4). Indeed, the minor allele of this SNP that we identified here as having more positive yields in polycultures compared to expected was also associated with earlier flowering at 16°C among the 60 genotypes we used from [37] (t-test, t= −2.6687, p = 0.0279). As the frequency of this early-flowering minor allele increased and the stand variance at this locus increased, DME also became more positive (Figure 4). We found the 26 flowering time candidate genes had a number of SNPs with nominally significant associations with biomass and DME in both treatments. Notably, SPL4 (AT1G53160), HEN2 (AT2G06990), ULP1B (AT4G00690), and ATH1 (AT4G32980) were significantly associated with DME in low resource stands (SNP frequency and variance associations, with candidate SNP-wide FDR < 0.05, Table S7).

**Figure 3.**
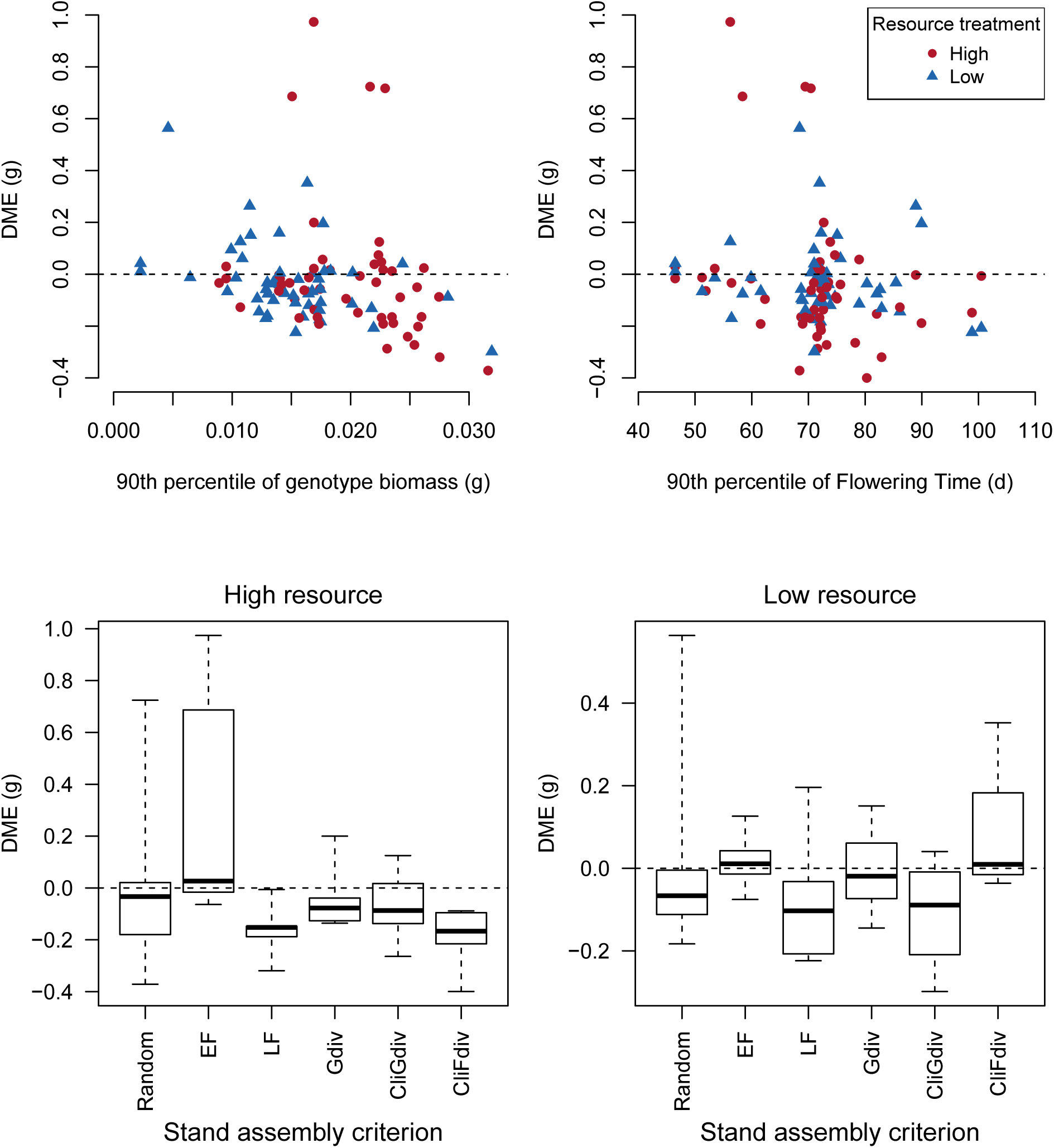
Polyculture deviations from monoculture expectation (DME) in comparison with measures of stand composition (top panels) or stand assembly criteria (bottom panels). (Top panels) Stands with large biomass genotypes, or late flowering genotypes (90^th^ percentiles) also tend to show lower DME (under-yielding), even more so than when measured via mean monoculture biomass or flowering time (Table S5). (Bottom panels) Stands assembled via different criteria show different DME. In particular, stands with genotypes predicted to be more early flowering (EF) had higher DME (over-yielding) compared to late flowering (LF) stands (Tukey HSD, FDR adjusted p = 0.0292). Gdiv=genomic diversity, CliGdiv=similar climate to study site but genomic diversity, CliFdiv=similar climate of origin to study site but flowering time diversity.

**Figure 4:**
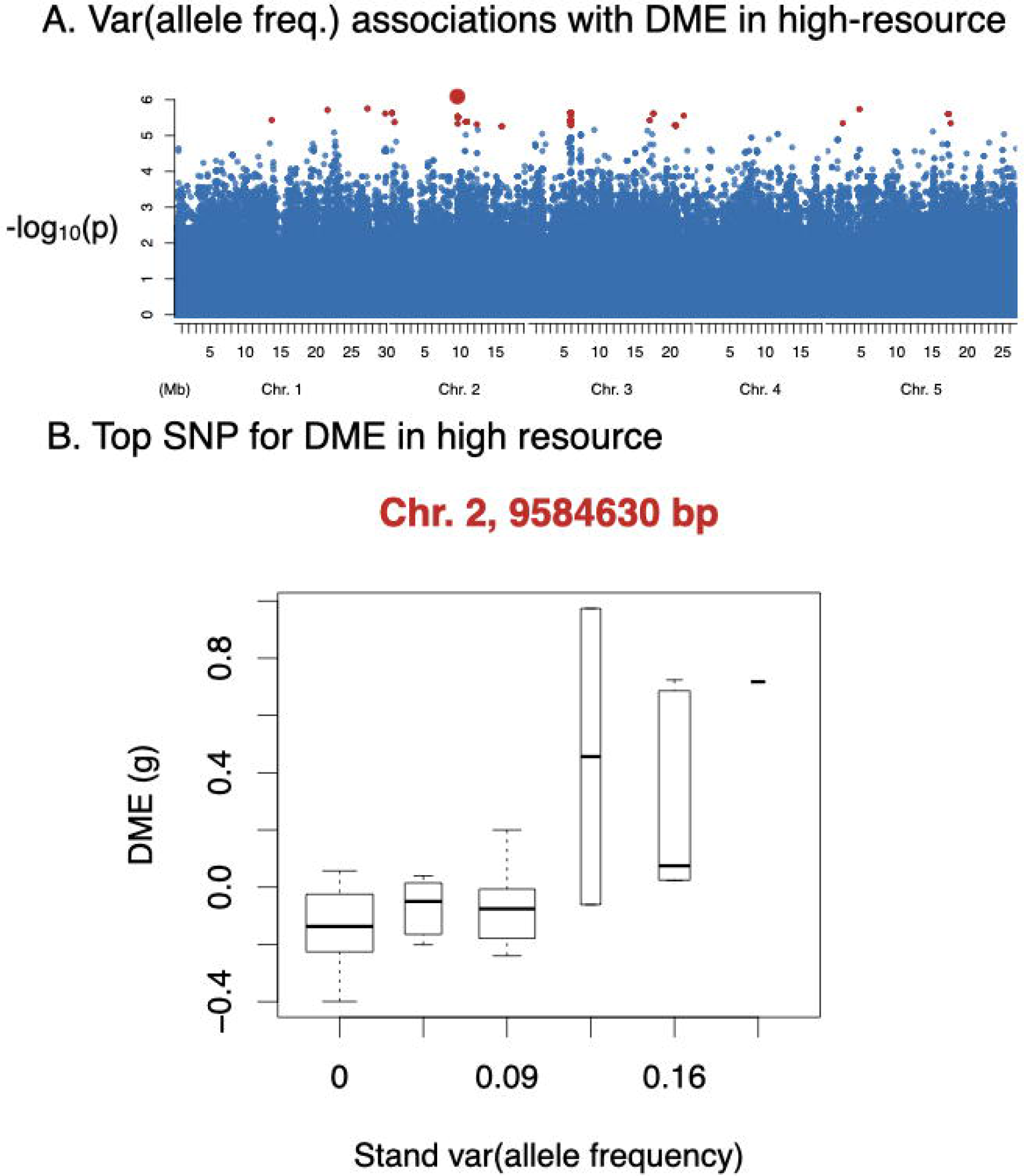
Associations between DME in the high resource treatment with var(allele frequency) across the genome (A) and at the top associated SNP (B). (A) Linear mixed model genome-wide association with 1.9M SNPs allele frequency variance (var = *pq*) versus deviation from monoculture expectation (DME) in high resource environments. SNPs significant with FDR < 0.1 are shown in red. The top SNP is highlighted with a large red dot in panel A and is featured in panel B (this was also the top SNP for allele frequency associations with DME). Boxplot (panel B) shows variance in allele frequency within polycultures (x-axis) at this SNP (chromosome 2, 9584630 bp) associated with DME of biomass (panel B y-axis) in the high resource environment.

## DISCUSSION

Diversity can affect the group-level properties of organisms, including ecosystem functions like carbon cycling and population trajectories. Diversity effects have been often studied via observation of uncontrolled systems that vary in species or genetic diversity [29,46]. In particular, species and higher levels of diversity have been demonstrated to have a range of effects on ecosystem function via multiple mechanisms [8,47]. We built on this research to test how intraspecific variation in genomic and trait composition and diversity affect the performance of an established non-native species in the field. We found that the diversity of genotypes did not increase biomass production; instead polycultures showed lower biomass production than expected, suggesting interference competition. This low polyculture yield was associated with stands containing late-flowering genotypes that had high productivity in monoculture. We identified genetic variation in a known flowering time locus associated with these effects. We also found a weak but significant increase in fecundity with the number of genotypes in a treatment, for unclear reasons.

### Diversity effects on average individual fecundity

Because of logistical and (in the case of invasive species) bioethical constraints, many studies of the effect of diversity on population performance focus on the impact on biomass production and may not assess fecundity directly (see [2,19,20]). While none of the diversity metrics we tested had an effect on biomass production, we did find a small but significant effect of genotype number on stand level fecundity in the high resource environment (Fig 2c). The mechanism underlying this effect is unclear, given the lack of an effect on biomass. This may be due to sampling effects, specifically where greater diversity increases the chance a group will contain early flowering and more fecund genotypes. Alternately, interference competition may trigger earlier reproduction in small (especially early flowering) genotypes to a greater degree than large (especially late flowering) genotypes, leading to positive diversity-fecundity associations.

### Diversity effects on stand-level biomass

We found no evidence that increased genetic diversity led to greater stand-level aboveground biomass production of Arabidopsis in the field, highlighting the complexity of stand-level properties in even this model plant. Our findings may be explained by a lack of such genetic diversity effects in Arabidopsis populations in general, or by technical/logistical constraints that made us miss such effects. First, diversity effects might not have been observed if positive effects of diversity (complementarity of resource use or pathogen susceptibility) were opposed by negative effects (interference competition) [7] or if diversity effects did not exist. Our finding of reduced polyculture yield compared to expectation from monoculture (DME) suggests interference competition as one potential explanation. Second, diversity effects might have been overlooked if they occurred along dimensions that were not measured, such as seed survival or nutrient use [26]. Diversity effects might also have been overlooked if they occur across environmental gradients unobserved in our single year, two resource level, study [48,49].

By contrast, we found consistent under-yielding of polycultures of genotypes compared to the expectations from their biomass in monoculture and relative representation in the polycultures (deviation from monoculture expectation, DME, also known as “diversity effects”) [4,7]. Generally, negative DMEs are thought to arise from interference competition [7]. This process could play out in our experiment as follows: when grown with genotypes having different trait values, individuals exhibit reduced performance compared to in monoculture with genetically identical individuals, and apparently without much benefit to the performance of the higher performing genotypes. For example, [50] found that *Pseudomonas flourescens* genotypic diversity lead to reduced ecosystem function (host plant protection) due to antagonistic interference (mutual poisoning) among *Pseudomonas* genotypes. Similar patterns have been reported in plants, where individuals have higher performance among similar genotypes and reduced performance among different genotypes [24,27]. For example, [24] found reduced variance in biomass among individuals and higher individual fecundity of *Plantago lanceolata* when grown among related individuals, which he interpreted as being due to reduced interference competition in high relatedness stands (though the term “interference competition” was not used). This might occur if higher performing genotypes reduce access to resources of lower performing genotypes to a greater degree than those high performing genotypes benefit by being surrounded by fewer of their own type. Alternatively, higher performing genotypes in monoculture might be more limited by lower performing genotypes, with little benefit to lower performing genotypes. We found that polyculture plots containing later flowering and larger biomass (in monoculture) genotypes were those that had the strongest negative DME, possibly due to shading of different and early flowering smaller neighbors. This suggests that late-flowering genotypes may have been more likely to be involved in interference competition when resources were abundant, perhaps because these larger individual plants limited their competitors from resource capture (e.g. by shading) more than they benefited, reducing the overall total biomass [24]. Evidence suggests that size asymmetries leading to shading [51] as well as allelopathy against competitors or other trophic levels [52] can also lead to such interference competition. Other authors have sometimes interpreted these patterns as evidence of kin selection [53,54]. The interference competition explanation for negative DME seems more likely than plants’ ability to first recognize kin *per se* and then reduce interference with related neighbors (cf. [54]).

We demonstrated how genomic data on individuals in experimental stands can be used to identify genetic loci potentially affecting group-level performance, although we did not identify effects of whole genome diversity on stand performance. Recent studies using Arabidopsis grown in pots, combined with linkage [23] or association mapping [35] have demonstrated complementary genomic approaches using a smaller number of genotypes and individuals per stand (two genotypes and three-four individuals/pot). Controlled experiments like these and our present study, combined with group-level performance and genetic data, could be applied to nearly any system. It is not clear whether such approaches could detect genetic loci operating in natural uncontrolled systems, where uncontrolled environmental variation plays an important but often unmeasured role in driving spatial variation in group-level performance. Here, we implemented a study of association of stand level performance with both allele frequency and variance in stands. However, note that to distinguish between effects of variance (diversity *per se*) versus frequency (composition) at a biallelic SNP, a given SNP would need to span a wide range of allele frequencies among stands. This condition is not likely met for most SNPs given they have low minor allele frequency, thus allele frequency and variance show a large degree of covariation, limiting our ability to distinguish the effects of diversity versus allelic composition at most loci.

### Previous studies on Arabidopsis

Previous studies in the laboratory have shown some evidence that certain combinations of Arabidopsis genotypes may result in higher stand level performance. In greenhouse populations of Arabidopsis, both additive and non-additive benefits of up to eight genotypes per stand (from a pool of 23) were found to increase emergence, biomass, flowering duration, and fecundity [19]. [23] used an Arabidopsis biparental mapping population to identify a single locus causing overyielding when grown in diversity in pots. Additionally, [35] used pot experiments on pairs of genotypes in a greenhouse, combined with association mapping on an individual genotype’s association with over-versus under-yielding. Our study was conducted in the field where plants were subject to a wide range of potential stressors. It is noteworthy that [55] found that natural stands in Germany with a single genotype tended to have fewer individuals than stands with multiple genotypes, suggesting potential positive diversity effects in nature.

### A priori prediction of stand-level performance

Ecology has been criticized for not being a predictive science. Here we tested our knowledge of the system with *a priori* sets of stands predicted to have divergent performance. We found that our predictions were most closely associated with stand-level yield driven by flowering time differences among genotypes. Going forward, if we were to construct a productive Arabidopsis stand with high fecundity individuals at our site, we would choose a set of multiple distinct genotypes, that are late flowering, and with low SLA (but note some late-flowering genotypes did not flower here, and this strategy may only work in certain environments).

It is no accident that we chose flowering time as the one trait to include in our prediction criteria. Flowering time has well-documented links to a major axis of life history and physiological variation [39,56], and is so intensely studied for agronomic interest. The use of phenotyped and genotyped individuals in such experiments are a way forward for predictive approaches to group performance and diversity.

## Conclusions

Biodiversity can affect the properties of groups of organisms and may be important to understanding the performance and persistence of both natural and introduced populations and communities. Here we did not identify strong effects of genetic diversity *per se* on the performance of stands of non-native Arabidopsis populations but we found a pattern of higher biomass yield when genetic composition was shifted to late-flowering stands. We also identified that in polyculture, the presence of later-flowering genotypes led to lower-than expected yield, possibly due to interference competition. Our research here on stand-level effects on performance in the field for the model plant Arabidopsis suggest a way forward to understanding the importance of such effects in nature.

## Supporting information

MainFieldExperimentalData

## Acknowledgments

Connor Campana, Timothy Gilpatrick, Sarah Lucas, and Ken Hull assisted with fieldwork. Victoria DeLeo, Emily Bellis, Lua Lopez, and Jonathan Kizer assisted with planting. Paras Patel and Crosley Williams assisted with sorting seeds and planting. Leland Burghard assisted with plant care. Megan Vahsen provided useful discussions on experimental design. This work was supported by a Pennsylvania State University Eberly Postdoctoral Fellowship and NSF Idaho EPSCoR Program award OIA-1757324 to KGT. We lastly thank the cattle that trampled only small part of our experiment.

