## Supplemental methods, tables, figures for "Effects of genomic and functional diversity on stand-level productivity and performance of non-native Arabidopsis"

**Online supplement**

**Methods S1**

*Seeds used in experiments*

The genotypes used in experiments were nearly evenly distributed among 7 genetic clusters or admixed genotypes identified by [37] (though we did not have any Swedish genotypes). We excluded any genotypes that were reported as contaminants by the supplying lab [1]. Seeds were obtained from the Arabidopsis Biological Resource Center and were bulked November 2016 through June 2017 in a growth room at a constant temperature between 20-25 °C, under 90-120 PAR with a 16-hour photoperiod.

*Assembly of prediction stands*

We used five different criteria to generate these 'prediction stands': (1) genomic diversity among accessions, (2) climate similarity of collection location to study site combined with genomic diversity, (3) early flowering genotypes, (4) late flowering genotypes, (5) climate similarity of collection location to our study site, combined with flowering time diversity. For (1) and (2), we sought to maximize genome-wide diversity. We began with a random genotype and then followed adding genotypes based on lowest average kinship with already selected genotypes (kinship matrix calculated by [2], based on whole genome resequencing data). For (2), we conducted the same exercise except that we began with the ecotype closest in climate space to the study site (see below) and then selected a subset of ecotypes closest in climate space to the study site. From this subset we then chose the least closely related genotype (lowest kinship coefficient) and then followed adding genotypes based on the combined ranks of a) low average kinship with already chosen genotypes and b) low climate distance from the experimental site.

For (2) and (3), we calculated principal components of climate for the 60 ecotype locations of origin combined with our experimental site, using variables potentially important in local adaptation in Arabidopsis [3–9]: mean April temperature, mean diurnal temperature range, minimum temperature of coldest month, mean monthly minimum and mean monthly maximum temperatures during the growing season, growing season precipitation relative to potential evapotranspiration, inter-annual coefficient of variation in growing season precipitation, growing season length, coefficient of variation in monthly precipitation across the years, and coefficient of variation in monthly precipitation during the growing season [8,10–12]. The first five PCs accounted for 93% of the total climate variation. The first PC largely corresponded to an axis of temperature variation among sites. The second PC largely corresponded to high precipitation variability, high maximum growing season temperatures, a large diurnal temperature range, and dry growing seasons at one extreme. We then calculated each ecotype's Manhattan distance from the experimental site in the space of the first five principal components.

For (3-5) we calculated average flowering time for these genotypes grown at 10ºC and 16ºC in growth chambers, using published data [2]. For (3) early flowering stands, we selected the lower tail of flowering times (tail size depending on number of genotypes in stand: 0.1, 0.3, or 0.5) and then randomly selected genotypes within that tail. For (4) the late flowering stands we used the same method with similarly sized upper tails of the flowering time distribution. For (5), we again began with a genotype from similar climate to the study site and then selected a subset of genotypes closest in climate space to the study site. From this subset we chose the genotype with the most different flowering time and then followed adding genotypes based on combined ranks of low flowering time similarity with already chosen genotypes and low climate distance from the experimental site.

*Randomly assembled stands*

In assembling random stands we sought to limit similarity of stands within a treatment and within stands having the same number of genotypes [19,40]. Pairwise comparisons were made for all stand compositions (including prediction stands) to minimize redundancy across stands. If a potential random stand genotype composition overlapped by more than the allowed number of genotypes with a prediction stand or a previously accepted random stand (no repeat lines allowed among 2-genotype stands, 4 repeat lines allowed among 10-genotype stands, 9 repeat lines allowed among 20-genotype stands), that stand composition was rejected and a new one randomly generated. We generated random polycultures of either 2, 10, or 20 different randomly selected genotypes, with ten stand replicates per diversity level, for a total of 30 different randomly generated polycultures. All genotypes were included within the 10 stand replicates, randomly assigned to replicates such that each replicate had a different assortment of genotypes but had the same diversity level. In the resulting final stand compositions (including predicted and random stands), each 2-genotype stand was unique; each pair of 10-genotype stands shared 0.795 genotypes on average; and each pair of 20-genotype stands shared 1.725 genotypes on average. The average genotype was represented in 0.67 (SD=0.46) 2-genotype stands, 3.33 (SD=2.08) 10-genotype stands, and 6.67 (SD=2.26) 20-genotype stands.

*Experimental conditions*

All seeds were stratified on April 30, 2018 in tap water for 4 days and planted on May 3-4, 2018 in square 10.16 x 10.16 x 8.89 cm cow manure pots (CowPots, East Canaan, CT), each of which contained a 20-plant stand. Assigned genotypes for each stand were randomly planted into a gridded location within the stand (4 rows of 5 plants each) and were planted approximately 1.5 cm apart. Stands designated as a low resource treatment were filled with 50% medium commercial grade sand and 50% potting mix (Premier pro-mix PGX), whereas stands representing a high resource treatment contained 100% potting mix. All potting components were autoclaved prior to filling pots.

After each stand was planted, it was placed into darkness at 4°C for at least 24 hours. Once all stands were planted, stands were placed into a growth room at 20-25°C with a 20 hour photoperiod with light levels between 100-200 PAR, and covered until germination. Once cotyledons were expanded, plants were thinned to one plant per position in stand (giving 20 plants), and trays of pots were rotated every other day between the growth room and a Percival growth chamber (Model PGC-40L2, Percival Scientific, Inc., Perry, IA) set to 600-620 PAR and daily temperature cycles from 15 to 26°C, to acclimate plants to higher light levels and cooler night temperatures. After thinning and eight days prior to transplanting into the field, the germination rate for each planting position within stands was 92%.

Pots were transplanted to the 850 $m^2$ ploughed field site on May 24, 2018. The site was located at 314 m asl in a meadow on flat topography at the bottom of a slope, with soils that were high in clay (Hagerstown silty clay loam, USDA Web Soil Survey https://websoilsurvey.nrcs.usda.gov/app/WebSoilSurvey.aspx). Pot locations in the field were randomly assigned and pots were buried into the soil so that the pot soil was at the same level as the field soil. All monoculture, polyculture, and prediction pots (stands) across both treatments were randomly assigned locations within the field plot (Figure S2). Stands were organized into four blocks, each block containing 120 stands (4 x 30 stands). There was a 15.24 cm buffer between stands within a block, and a 91.44 cm buffer between blocks and between blocks and the edge of the ploughed site. The field site was completely fenced in and was weeded for non-Arabidopsis plants twice weekly. Initial plant number per stand was assessed by counting the surviving seedlings five days after transplanting stands into the field, totaling 8384 plants across the 480 stands (averaging 17.5 plants/stand on May 29).

**Table S1.** Results of linear mixed models of plastic trait response to soil treatments in monocultures.

| *Trait* | *High - low resource trait change* | *SE* | *t Value* | *p* |
|---|---|---|---|---|
| log(SLA) (ln (cm2/mg)) | -0.1035 | 0.0404 | -2.561 | 0.0626 |
| Date of first flowering (days) | -0.2943 | 1.1325 | -0.26 | 0.8078 |
| Date of first flowering, including plants that never flowered (days) | -1.354 | 0.6628 | -2.043 | 0.1105 |
| AGB per plant (g) | 0.0035 | 0.0012 | 3.042 | 0.0383 |
| Fecundity per reproductive plant (mm fruit length) | 21.82 | 5.93 | 3.676 | 0.021 |

**Table S2**. Broad-sense heritabilities.

| *Trait* | *High resource* | *Low resource* |
|---|---|---|
| log(SLA) | 0.42 | 0.52 |
| Date of first flowering | 0.88 | 0.80 |
| Date of first flowering, including plants that never flowered | 0.87 | 0.92 |
| Date of 50% flowering | 0.73 | 0.54 |
| AGB per plant | 0.43 | 0.41 |
| Fecundity per reproductive plant | 0.28 | 0.67 |

**Table S3**. Linear models of plot-level fecundity of Arabidopsis stands measured in the field common garden.

| **Covariate** | **Treatment** | **Estimate** | p | $R^2_{adjusted}$ |
|---|---|---|---|---|
| *Diversity metrics* | | | | |
| Diversity Level | High | 2.20762 | 0.0289* | 0.046 |
| Diversity Level | Low | 0.63101 | 0.5733 | |
| Mean genetic distance | High | 26.37127 | 0.0944 | 0.035 |
| Mean genetic distance | Low | -2.10719 | 0.9026 | |
| *Composition metrics* | | | | |
| Distance in PCs 1-5 (mean) | High | 1.52930 | 0.5467 | 0.023 |
| Distance in PCs 1-5 (mean) | Low | 0.16076 | 0.9483 | |
| SLA breeding value (mean) | High | -30.27388 | 0.3107 | 0.017 |
| SLA breeding value (mean) | Low | -3.20336 | 0.9215 | |
| FT (mean) | High | -0.05192 | 0.9461 | 0.038 |
| FT (mean) | Low | 1.43961 | 0.0625 | |

Plot level fecundity measured as estimated mean seed number (described in text). Asterisk indicates significance of covariate in linear models at $P$ <0.05.

**Table S4.** Linear models of per plant biomass of Arabidopsis stands measured in the field common garden.

| Covariate | Treatment | Estimate | p | $R^2_{adjusted}$ |
|---|---|---|---|---|
| *Diversity metrics* | | | | |
| Diversity Level | High | -0.00004 | 0.7958 | 0.015 |
| Diversity Level | Low | -0.00011 | 0.4743 | |
| Mean genetic distance | High | -0.00161 | 0.4251 | 0.017 |
| Mean genetic distance | Low | -0.00192 | 0.3577 | |
| *Composition metrics* | | | | |
| Distance in PCs 1-5 (mean) | High | -0.00049 | 0.1298 | 0.022 |
| Distance in PCs 1-5 (mean) | Low | -0.00031 | 0.3325 | |
| SLA breeding value (mean) | High | -0.00926 | 0.0095* | 0.029 |
| SLA breeding value (mean) | Low | -0.00392 | 0.3388 | |
| FT (mean) | High | 0.00013 | 0.0204* | 0.050 |
| FT (mean) | Low | 0.00017 | 0.0024* | |

Plot level biomass measured as aboveground dry weight at harvest (described in text). Asterisk indicates significance of covariate in linear models at $P < 0.05$.

**Table S5**. Linear models of per plant canopy area of Arabidopsis stands measured in the field common garden.

| **Covariate** | **Treatment** | **Estimate** | p | $R^2_{adjusted}$ |
|---|---|---|---|---|
| *Diversity metrics* | | | | |
| Diversity Level | High | -0.00121 | 0.7710 | 0.201 |
| Diversity Level | Low | 0.00009 | 0.9831 | |
| Mean genetic distance | High | -0.02716 | 0.6574 | 0.202 |
| Mean genetic distance | Low | -0.02649 | 0.6648 | |
| *Composition metrics* | | | | |
| Distance in PCs 1-5 (mean) | High | -0.00051 | 0.9572 | 0.201 |
| Distance in PCs 1-5 (mean) | Low | -0.00053 | 0.9539 | |
| SLA breeding value (mean) | High | -0.02102 | 0.8549 | 0.205 |
| SLA breeding value (mean) | Low | -0.03843 | 0.7463 | |
| FT (mean) | High | -0.00075 | 0.6662 | 0.202 |
| FT (mean) | Low | -0.00026 | 0.8782 | |

**Table S6.** Linear model statistics for associations with DME (deviation from monoculture expectation)

| Covariate | Treatment | Estimate | p | $R^2_{adjusted}$ |
|---|---|---|---|---|
| *Diversity metrics* | | | | |
| Mean genome-wide similarity | High | -0.04639 | 0.7244 | <0 |
| Mean genome-wide similarity | Low | 0.02015 | 0.8821 | |
| NJ tree branch length | High | 0.00108 | 0.6369 | <0 |
| NJ tree branch length | Low | -0.00028 | 0.9063 | |
| N genotypes | High | 0.00330 | 0.3896 | <0 |
| N genotypes | Low | -0.00024 | 0.9526 | |
| *Composition metrics* | | | | |
| FT (90th percentile) | High | -0.00514 | 0.0170 | 0.0275 |
| FT (90th percentile) | Low | -0.00060 | 0.7883 | |
| FT (mean) | High | -0.00643 | 0.0167 | 0.0290 |
| FT (mean) | Low | -0.00121 | 0.6532 | |
| AGB (90th percentile) | High | -11.62425 | 0.0030 | 0.0954 |
| AGB (90th percentile) | Low | -11.58907 | 0.0309 | |
| AGB (median) | High | -14.68487 | 0.0133 | 0.0812 |
| AGB (median) | Low | -18.78808 | 0.0167 | |
| SLA (mean) | High | 0.70971 | 0.3544 | <0 |
| SLA (mean) | Low | 0.47972 | 0.3066 | |
| Distance in PCs 1-5 env (mean) | High | 0.03969 | 0.0485 | 0.0096 |
| Distance in PCs 1-5 env (mean) | Low | 0.00313 | 0.8785 | |

**Table S7**. Biomass and DME associations for SNPs within 3kb of 26 candidate flowering time genes. These genes were identified as candidates for non-vernalized flowering time by [13]. False discovery rate (FDR) control was applied to all of these candidate SNPs for a given response variable (i.e. combination of treatment, response variable, and covariate), yielding *q* values (see attached file).

**Figure S1.** Map of location of origin of each of the 60 native-range genotypes used in this study (blue circles).

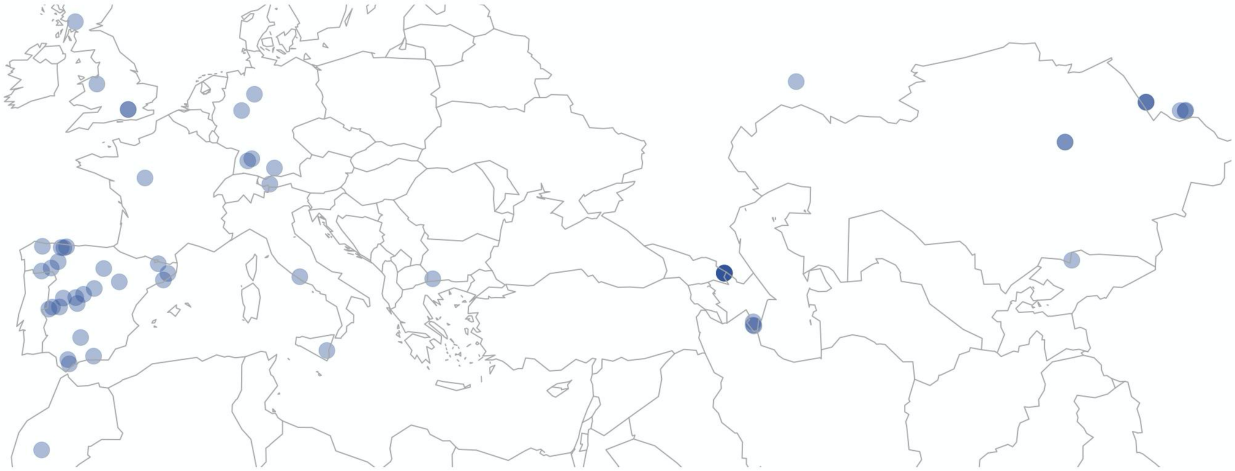

**Figure S2.** Field plot design with four blocks. Each square represents a stand. Total plot area was 85 square meters (10 x 8.5 meters).

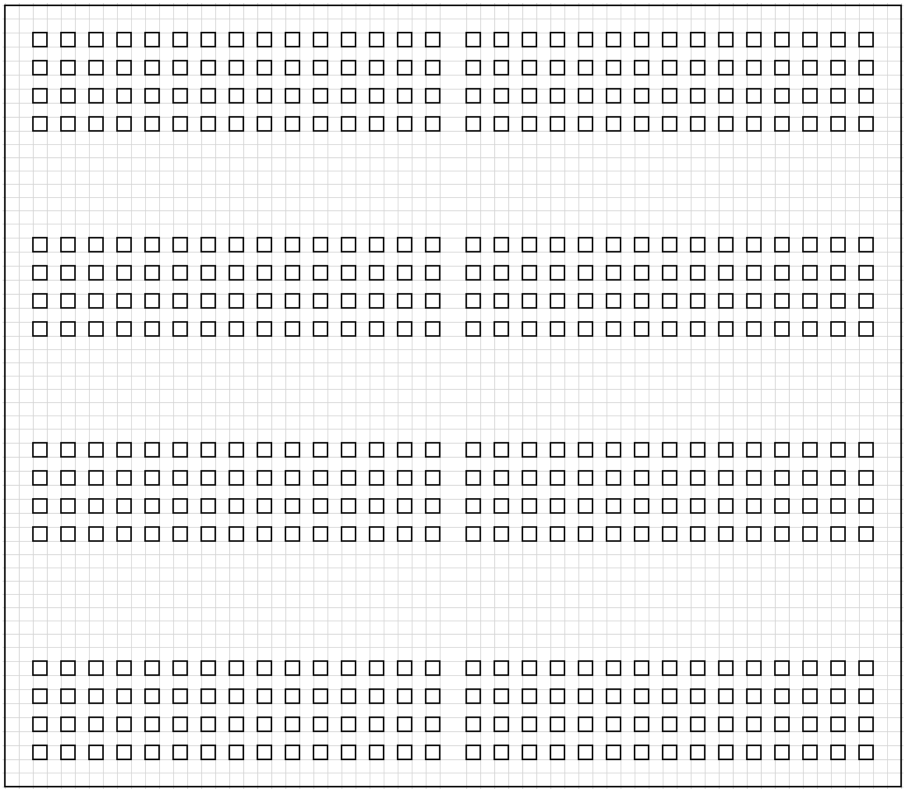

**Figure S3**. Percent volumetric water content (VWC%) for each treatment in addition to the surrounding high clay soil was monitored each week within each block using EM-50 dataloggers and EC-5 probes (METER Group Inc., Pullman, WA). At the beginning of the experiment (June 8, 2018), mean VWC for the high resource stands was 13.85%, 9.9% for the low resource stands, and 29.25% for the surrounding field soil (which had high clay content). Toward the end of the experiment (July 12, 2018), VWC for the high resource stands was 11.95%, 5.38% for the low resource stands, and 26.78% for the surrounding field soil.

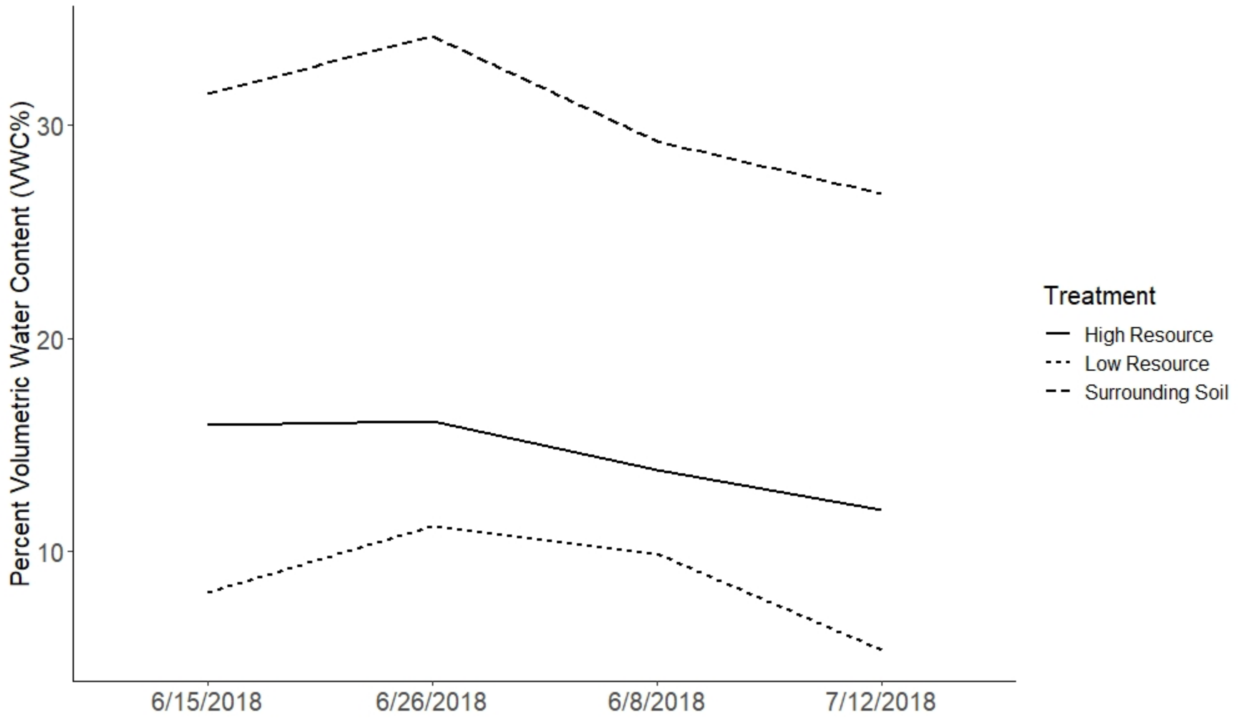

**Figure S4**. Genome-wide associations between stand biomass or DME and allele frequency within stands, or variance among individuals within stands. y-axes show -$\log_{10}$ p-values of association, the five chromosomes of Arabidopsis are arrayed on the x-axes.

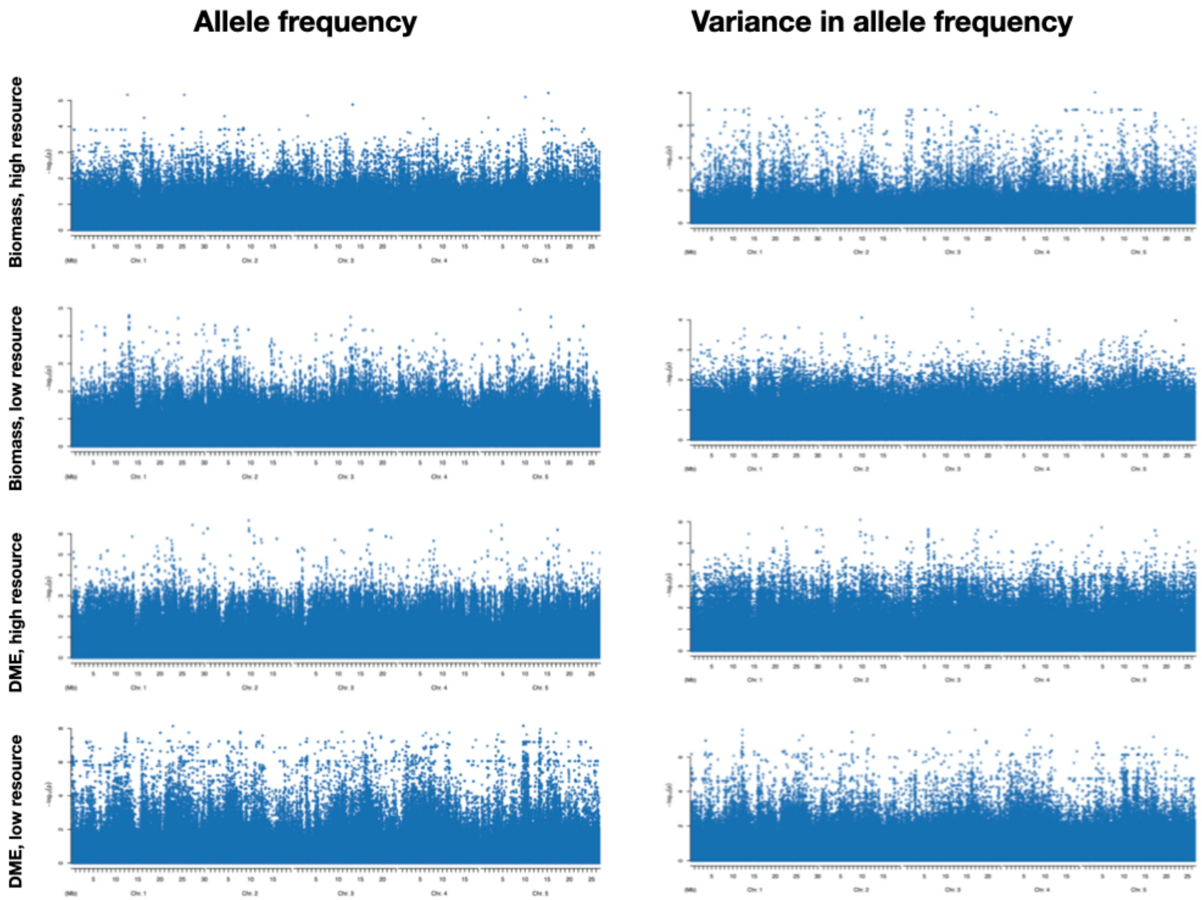

**Figure S5**. Biomass and fecundity performance of prediction stands in high and low resource environments. Performance of prediction plots and random polyculture plots in high and low resource environments. Criteria for prediction plots described in text. ‘EF’ = ‘early flowering’, 12 stands. ‘Gdiv’ = genetic diversity criterion, 11 stands. ‘LF’ = ‘late flowering’, 11 stands. ‘SimClimFdiv’ = similar climate, flowering time diversity criterion, 10 stands. ‘SimClimGdiv’ = similar climate, genetic diversity criterion, 11 stands. ‘div’ = all remaining randomly assembled polyculture stands, 55 stands. (a) Per plant biomass at harvest. (b) Plot level Estimated seed number. Because only two stands from the LF criteria produced fruit, it was not included in analyses of fecundity.

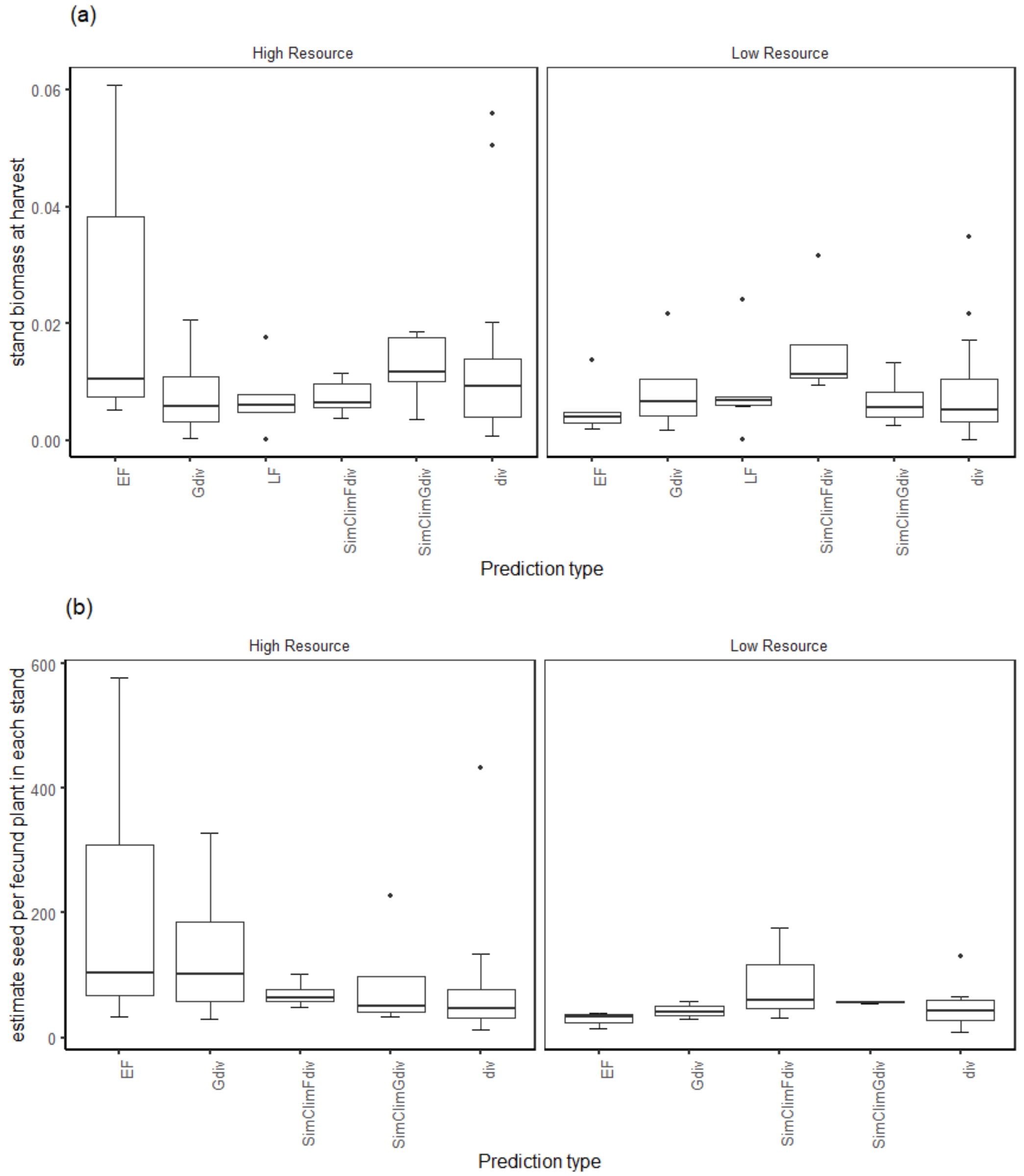

6. Gienapp P, Fior S, Guillaume F, Lasky JR, Sork VL, Csilléry K. 2017 Genomic quantitative genetics to study evolution in the wild. *Trends Ecol. Evol.*
